# Sampling networks of ecological interactions

**DOI:** 10.1101/025734

**Authors:** Pedro Jordano

## Abstract

1. Sampling ecological interactions presents similar challenges, problems, potential biases, and constraints as sampling individuals and species in biodiversity inventories. Interactions are just pairwise relationships among individuals of two different species, such as those among plants and their seed dispersers in frugivory interactions or those among plants and their pollinators. Sampling interactions is a fundamental step to build robustly estimated interaction networks, yet few analyses have attempted a formal approach to their sampling protocols.
2. Robust estimates of the actual number of interactions (links) within diversified ecological networks require adequate sampling effort that needs to be explicitly gauged. Yet we still lack a sampling theory explicitly focusing on ecological interactions.
3. While the complete inventory of interactions is likely impossible, a robust characterization of its main patterns and metrics is probably realistic. We must acknowledge that a sizable fraction of the maximum number of interactions *I*_*max*_ among, say, *A* animal species and *P* plant species (i.e., *I*_*max*_ = *AP*) is impossible to record due to forbidden links, i.e., life-history restrictions. Thus, the number of observed interactions *I* in robustly sampled networks is typically *I ≪ I_max_*, resulting in extremely sparse interaction matrices with low connectance.
4. Reasons for forbidden links are multiple but mainly stem from spatial and temporal uncoupling, size mismatches, and intrinsically low probabilities of interspecific encounter for most potential interactions of partner species. Adequately assessing the completeness of a network of ecological interactions thus needs knowledge of the natural history details embedded, so that forbidden links can be “discounted” when addressing sampling effort.
5. Here I provide a review and outline a conceptual framework for interaction sampling by building an explicit analogue to individuals and species sampling, thus extending diversity-monitoring approaches to the characterization of complex networks of ecological interactions. This is crucial to assess the fast-paced and devastating effects of defaunation-driven loss of key ecological interactions and the services they provide and the analogous losses related to interaction gains due to invasive species and biotic homogenization.

## Introduction

~~~
Biodiversity sampling is a labour-intensive activity, and sampling is often not sufficient to detect all or even most of the species present in an assemblage. Gotelli & Colwell (2011).
~~~

Biodiversity species assessment aims at sampling individuals in collections and determining the number of species represented. Given that, by definition, samples are incomplete, these collections do not enumerate the species actually present. The ecological literature dealing with robust estimators of species richness and diversity in collections of individuals is immense, and a number of useful approaches have been used to obtain such estimates (Magurran, 1988; Gotelli & Colwell, 2001; Colwell, Mao & Chang, 2004; Hortal, Borges & Gaspar, 2006; Colwell, 2009; Gotelli & Colwell, 2011; Chao *et al.*, 2014). Recent effort has been also focused at defining essential biodiversity variables (EBV) (Pereira *et al.*, 2013) that can be sampled and measured repeatedly to complement biodiversity estimates. Yet sampling species or taxa-specific EBVs is just probing a single component of biodiversity; interactions among species are another fundamental component, one that supports the existence, but in some cases also the extinction, of species. For example, the extinction of interactions represents a dramatic loss of biodiversity because it entails the loss of fundamental ecological functions (Valiente-Banuet *et al.*, 2014). This missed component of biodiversity loss, the extinction of ecological interactions, very often accompanies, or even precedes, species disappearance. Interactions among species are a key component of biodiversity and here we aim to show that most problems associated with sampling interactions in natural communities relate to problems associated with sampling species diversity, even worse. We consider pairwise interactions among species at the habitat level, in the context of alpha diversity and the estimation of local interaction richness from sampling data (Chao *et al.*, 2014). In the first part we provide a succinct overview of previous work addressing sampling issues for ecological interaction networks. In the second part, after a short overview of asymptotic diversity estimates (Gotelli & Colwell, 2001), we discuss specific rationales for sampling the biodiversity of ecological interactions. Most of the examples come from the analysis of plant-animal interaction networks, yet are applicable to other types of species-species interactions.

Interactions can be a much better indicator of the richness and diversity of ecosystem functions than a simple list of taxa and their abundances and/or related biodiversity indicator variables (EBVs). Thus, sampling interactions should be a central issue when identifying and diagnosing ecosystem services (e.g., pollination, natural seeding by frugivores, etc.). Fortunately, the whole battery of biodiversity-related tools used by ecologists to sample biodiversity (species, *sensu stricto*) can be extended and applied to the sampling of interactions. Analogs are evident between these approaches (see Table 2 in Colwell, Mao & Chang, 2004). Monitoring interactions is a biodiversity sampling and is subject to similar methodological shortcomings, especially under-sampling (Jordano, 1987; Jordano, Vázquez & Bascompte, 2009; Coddington *et al.*, 2009; Vázquez, Chacoff & Cagnolo, 2009; Dorado *et al.*, 2011; Rivera-Hutinel *et al.*, 2012). For example, when we study mutualistic networks, our goal is to make an inventory of the distinct pairwise interactions that made up the network. We are interested in having a complete list of all the pairwise interactions among species (e.g., all the distinct, species-species interactions, or links, among the pollinators and flowering plants) that do actually exist in a given community. Sampling these interactions thus entails exactly the same problems, limitations, constraints, and potential biases as sampling individual organisms and species diversity. As Mao & Colwell (2005) put it, these are the workings of Preston’s demon, the moving “veil line” (Preston, 1948) between the detected and the undetected interactions as sample size increases.

Early efforts to recognize and solve sampling problems in analyses of interactions stem from research on food webs and to determine how undersampling biases food web metrics (Martinez, 1991; Cohen *et al.*, 1993; Martinez, 1993; Bersier, Banasek-Richter & Cattin, 2002; Brose, Martinez & Williams, 2003; Banasek-Richter, Cattin & Bersier, 2004; Wells & O’Hara, 2012). In addition, the myriad of classic natural history studies documenting animal diets, host-pathogen infection records, plant herbivory records, etc., represent efforts to document interactions occurring in nature. All of them share the problem of sampling incompleteness influencing the patterns and metrics reported. Yet, despite the early recognition that incomplete sampling may seriously bias the analysis of ecological networks (Jordano, 1987), only recent studies have explicitly acknowledged it and attempted to determine its influence (Ollerton & Cranmer, 2002; Nielsen & Bascompte, 2007; Vázquez, Chacoff & Cagnolo, 2009; Gibson *et al.*, 2011; Olesen *et al.*, 2011; Chacoff *et al.*, 2012; Rivera-Hutinel *et al.*, 2012; Olito & Fox, 2014; Bascompte & Jordano, 2014; Vizentin-Bugoni, Maruyama & Sazima, 2014; Frund, McCann & Williams, 2015). The sampling approaches have been extended to predict patterns of coextintions in interaction assemblages (e.g., hosts-parasites) (Colwell, Dunn & Harris, 2012). Most empirical studies provide no estimate of sampling effort, implicitly assuming that the reported network patterns and metrics are robust. Yet recent evidences point out that number of partner species detected, number of actual links, and some aggregate statistics describing network patterns, are prone to sampling bias (Nielsen & Bascompte, 2007; Dorado *et al.*, 2011; Olesen *et al.*, 2011; Chacoff *et al.*, 2012; Rivera-Hutinel *et al.*, 2012; Olito & Fox, 2014; Frund, McCann & Williams, 2015). Most of these evidences, however, come either from simulation studies (Frund, McCann & Williams, 2015) or from relatively species-poor assemblages. Most certainly, sampling limitations pervade biodiversity inventories in tropical areas (Coddington *et al.*, 2009) and we might rightly expect that frequent interactions may be over-represented and rare interactions may be missed entirely in studies of mega-diverse assemblages (Bascompte & Jordano, 2014); but, to what extent?

## Sampling interactions: methods

When we sample interactions in the field we record the presence of two species that interact in some way. For example, Snow and Snow (1988) recorded an interaction whenever they saw a bird “touching” a fruit on a plant. We observe and record feeding observations, visitation, occupancy, presence in pollen loads or in fecal samples, etc., of *individual* animals or plants and accumulate pairwise interactions, i.e., lists of species partners and the frequencies with which we observe them. Therefore, estimating the sampling completeness of pairwise interactions for a whole network, requires some gauging of how the number (richness) of distinct pairwise interactions accumulates as sampling effort is increased) and/or estimating the uncertainty around the missed links (Wells & O’Hara, 2012).

Most types of ecological interactions can be illustrated with bipartite graphs, with two or more distinct groups of interacting partners (Bascompte & Jordano, 2014); for illustration purposes I’ll focus more specifically on plant-animal interactions. Sampling interactions requires filling the cells of an interaction matrix with data. The matrix, Δ = *AP* (the adjacency matrix for the graph representation of the network), is a 2D inventory of the interactions among, say, *A* animal species (rows) and *P* plant species (columns) (Jordano, 1987; Bascompte & Jordano, 2014). The matrix entries illustrate the values of the pairwise interactions visualized in the Δ matrix, and can be 0 or 1, for presence-absence of a given pairwise interaction, or take a quantitative weight *w*_*ji*_ to represent the interaction intensity or unidirectional effect of species *j* on species *i* (Bascompte & Jordano, 2014; Vazquez *et al.*, 2015). The outcomes of most ecological interactions are dependent on frequency of encounters (e.g., visit rate of pollinators, number of records of ant defenders, frequency of seeds in fecal samples). Thus, a frequently used proxy for interaction intensities *w*_*ji*_ is just how frequent new interspecific encounters are, whether or not appropriately weighted to estimate interaction effectiveness (Vazquez, Morris & Jordano, 2005).

We need to define two basic steps in the sampling of interactions: 1) which type of interactions we sample; and 2) which type of record we get to document the existence of an interaction. In step #1 we need to take into account whether we are sampling the whole community of interactor species (all the animals, all the plants) or just a subset of them, i.e., a sub matrix Δ_*m,n*_ of *m < A* animal species and *n < P* plant species of the adjacency matrix Δ_*AP*_ (i.e., the matrix representation of interactions among the partner species). Subsets can be: a) all the potential plants interacting with a subset of the animals (Fig. 1a); b) all the potential animal species interacting with a subset of the plant species (Fig. 1b); c) a subset of all the potential animal species interacting with a subset of all the plant species (Fig. 1c). While some discussion has considered how to establish the limits of what represents a network (Strogatz, 2001) (in analogy to discussion on food-web limits; Cohen, 1978), it must be noted that situations a-c in Fig. 1 do not represent complete interaction networks. As vividly stated by Cohen *et al.* (1993): “*As more comprehensive, more detailed, more explicit webs become available, smaller, highly aggregated, incompletely described webs may progressively be dropped from analyses of web structure (though such webs may remain useful for other purposes, such as pedagogy)*”. Subnet sampling is generalized in studies of biological networks (e.g., protein interactions, gene regulation), yet it is important to recognize that most properties of subnetworks (even random subsamples) do not represent properties of whole networks (Stumpf, Wiuf & May, 2005).

**Figure 1.**
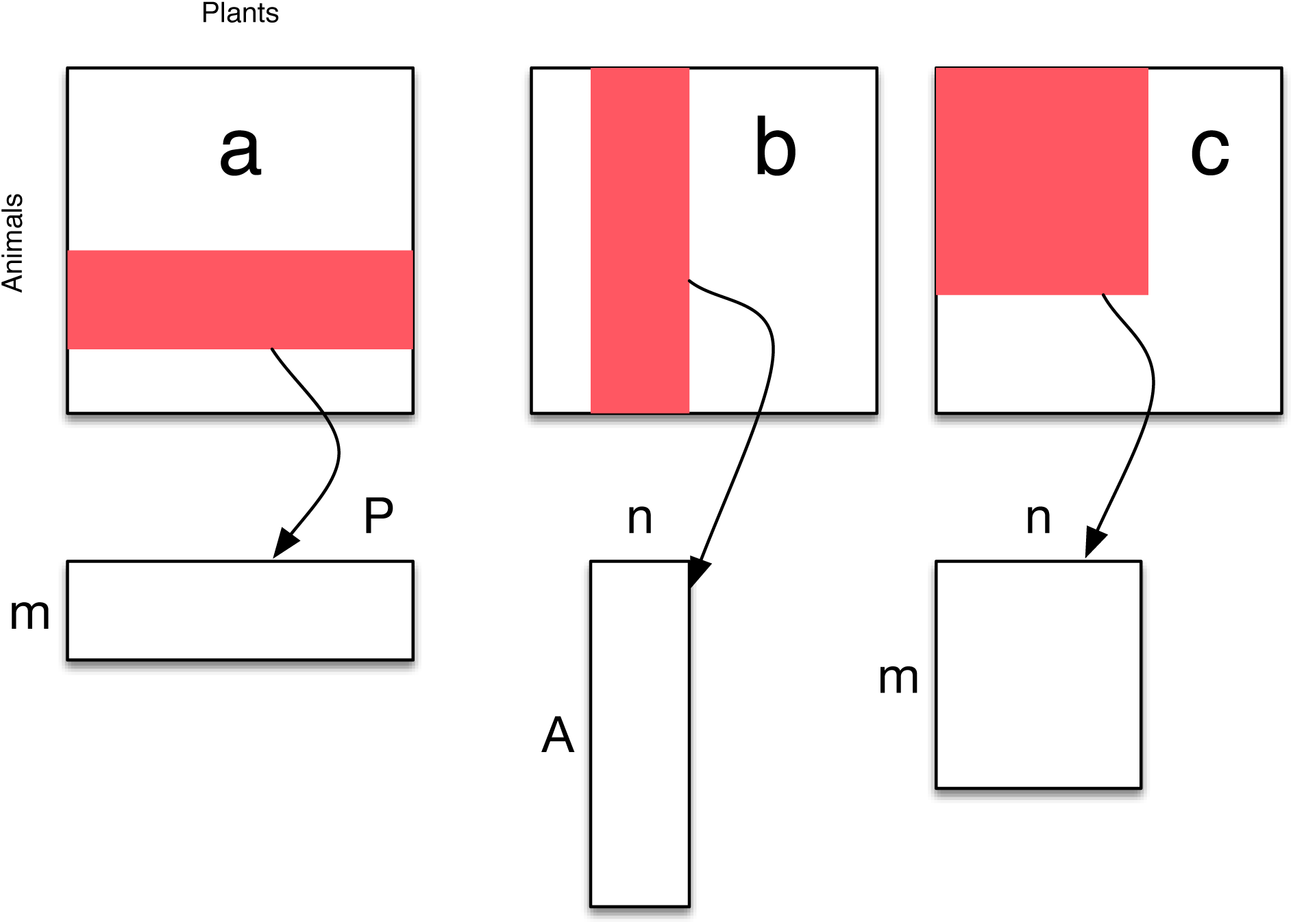
Sampling ecological interaction networks (e.g., plant-animal interactions) usually focus on different types of subsampling the full network, yielding submatrices Δ[*m, n*] of the full interaction matrix Δ with *A* and *P* animal and plant species. a) all the potential plants interacting with a subset of the animals (e.g., studying just the hummingbird-pollinated flower species in a community); b) all the potential animal species interacting with a subset of the plant species (e.g., studying the frugivore species feeding on figs *Ficus* in a community); and c) sampling a subset of all the potential animal species interacting with a subset of all the plant species (e.g., studying the plant-frugivore interactions of the rainforest understory).

In step #2 above we face the problem of the type of record we take to sample interactions. This is important because it defines whether we approach the problem of filling up the interaction matrix in a “zoo-centric” way or in a “phyto-centric” way. Zoo-centric studies directly sample animal activity and document the plants ‘touched’ by the animal. For example, analysis of pollen samples recovered from the body of pollinators, analysis of fecal samples of frugivores, radio-tracking data, etc. Phyto-centric studies take samples of focal individual plant species and document which animals ‘arrive’ or ‘touch’ the plants. Examples include focal watches of fruiting or flowering plants to record visitation by animals, raising insect herbivores from seed samples, identifying herbivory marks in samples of leaves, etc.

Most recent analyses of plant-animal interaction networks are phyto-centric; just 3.5% of available plant-pollinator (*N* = 58) or 36.6% plant-frugivore (*N* = 22) interaction datasets are zoo-centric (see Schleuning *et al.*, 2012). Moreover, most available datasets on host-parasite (parasitoid) or plant-herbivore interactions are “host-centric” or phyto-centric (e.g., Thébault & Fontaine, 2010; Morris *et al.*, 2013; Eklöf *et al.*, 2013). This may be related to a variety of causes, like preferred methodologies by researchers working with a particular group or system, logistic limitations, or inherent taxonomic focus of the research questions. A likely result of phyto-centric sampling would be adjacency matrices with large *A* : *P* ratios. In any case we don’t have a clear view of the potential biases that taxa-focused sampling may generate in observed network patterns, for example by generating consistently asymmetric interaction matrices (Dormann *et al.*, 2009). System symmetry has been suggested to influence estimations of generalization levels in plants and animals when measured as *I*_*A*_ and *I*_*P*_ (Elberling & Olesen, 1999); thus, differences in *I*_*A*_ and *I*_*P*_ between networks may arise from different *A* : *P* ratios rather than other ecological factors (Olesen & Jordano, 2002).

Reasonably complete analyses of interaction networks can be obtained when combining both phyto-centric and zoo-centric sampling. For example, Bosch *et al.* (2009) showed that the addition of pollen load data on top of focal-plant sampling of pollinators unveiled a significant number of interactions, resulting in important network structural changes. Connectance increased 1.43-fold, mean plant connectivity went from 18.5 to 26.4, and mean pollinator connectivity from 2.9 to 4.1; moreover, extreme specialist pollinator species (singletons in the adjacency matrix) decreased 0.6-fold. Olesen *et al.* (2011) identified pollen loads on sampled insects and added the new links to an observation-based visitation matrix, with an extra 5% of links representing the estimated number of missing links in the pollination network. The overlap between observational and pollen-load recorded links was only 33%, underscoring the value of combining methodological approaches. Zoocentric sampling has recently been extended with the use of DNA-barcoding, for example with plant-herbivore (Jurado-Rivera *et al.*, 2009), host-parasiotid (Wirta *et al.*, 2014), and plant-frugivore interactions (González-Varo, Arroyo & Jordano, 2014). For mutualistic networks we would expect that zoo-centric sampling could help unveiling interactions of the animals with rare plant species or for relatively common plants species which are difficult to sample by direct observation. Future methodological work may provide significant advances showing how mixing different sampling strategies strengthens the completeness of network data. These mixed strategies may combine, for instance, timed watches at focal plants, spot censuses along walked transects, pollen load or seed contents analyses, monitoring with camera traps, and DNA barcoding records. We might expect increased power of these mixed sampling approaches when combining different methods from both phyto- and zoo-centric perspectives (Bosch *et al.*, 2009; Blüthgen, 2010). Note also that the different methods could be applied in different combinations to the two distinct sets of species. However, there are no tested protocols and/or sampling designs for ecological interaction studies to suggest an optimum combination of approaches. Ideally, pilot studies would provide adequate information for each specific study setting.

## Sampling interactions: rationale

The number of distinct pairwise interactions that we can record in a landscape (an area of relatively homogeneous vegetation, analogous to the one we would use to monitor species diversity) is equivalent to the number of distinct classes in which we can classify the recorded encounters among *individuals* of two different species. Yet, individual-based interaction networks have been only recently studied (Dupont, Trøjelsgaard & Olesen, 2011; Wells & O’Hara, 2012). The most usual approach has been to pool indiviudal-based interaction data into species-based summaries, an approach that ignores the fact that only a fraction of individuals may actually interact given a per capita interaction effect (Wells & O’Hara, 2012). Wells & O’Hara (2012) illustrate the pros and cons of the approach. We walk in the forest and see a blackbird *T m* picking an ivy *Hh* fruit and ingesting it: we have a record for *T m - Hh* interaction. We keep advancing and record again a blackbird feeding on hawthorn *Cm* fruits so we record a *T m - Cm* interaction; as we advance we encounter another ivy plant and record a blackcap swallowing a fruit so we now have a new *Sa - Hh* interaction, and so on. At the end we have a series of classes (e.g., *Sa - Hh*, *T m - Hh*, *T m - Cm*, etc.), along with their observed frequencies. Bunge & Fitzpatrick (1993) provide an early review of the main aspects and approaches to estimate the number of distinct classes *C* in a sample of observations.

Our sampling above would have resulted in a vector *n* = [*n*_1_*…n_C_* ]*′* where *n*_*i*_ is the number of records in the *i*^*th*^ class. As stressed by Bunge & Fitzpatrick (1993), however, the *i*^*th*^ class would appear in the sample if and only if *n*_*i*_ > 0, and we don’t know *a priori* which *n*_*i*_ are zero. So, *n* is not observable. Rather, what we get is a vector *c* = [*c*_1_*…c_n_*]*′* where *c*_*j*_ is the number of classes represented *j* times in our sampling: *c*_1_ is the number of singletons (interactions recorded once), *c*_2_ is the number of twin pairs (interactions with just two records), *c*_3_ the number of triplets, etc. The problem thus turns to be estimating the number of distinct classes *C* from the vector of *c*_*j*_ values and the frequency of unobserved interactions (see “The real missing links” below).

More specifically, we usually obtain a type of reference sample (Chao *et al.*, 2014) for interactions: a series of replicated samples (e.g., observation days, 1h watches, etc.) with quantitative information, i.e., recording the number of instances of each interaction type on each day. This replicated abundance data, can be treated in three ways: 1) Abundance data within replicates: the counts of interactions, separately for each day; 2) Pooled abundance data: the counts of interactions, summed over all days (the most usual approach); and 3) Replicated incidence data: the number of days on which we recorded each interaction. Assuming a reasonable number of replicates, replicated incidence data is considered the most robust statistically, as it takes account of heterogeneity among days (Colwell, Mao & Chang, 2004; Colwell, Dunn & Harris, 2012; Chao *et al.*, 2014). Thus, both presence-absence and weighted information on interactions can be accommodated for this purpose.

## The species assemblage

When we consider an observed and recorded sample of interactions on a particular assemblage of *A*_*obs*_ and *P*_*obs*_ species (or a set of replicated samples) as a reference sample (Chao *et al.*, 2014) we may have three sources of undersampling error that are ignored by treating a reference sample as a true representation of the interactions in well-defined assemblage: 1) some animal species are actually present but not observed (zero abundance or incidence in the interactions in the reference sample), *A*_0_; 2) some plant species are actually present but not observed (zero abundance or incidence in the interactions in the reference sample), *P*_0_; 3) some unobserved links (the zeroes in the adjacency matrix, *U L*) may actually occur but not recorded. Thus a first problem is determining if *A*_*obs*_ and *P*_*obs*_ truly represent the actual species richness interacting in the assemblage. To this end we might use the replicated reference samples to estimate the true number of interacting animal *A*_*est*_ and plant *P*_*est*_ species as in traditional diversity estimation analysis (Chao *et al.*, 2014). If there are no uniques (species seen on only one day), then *A*_0_ and *P*_0_ will be zero, and we have *A*_*obs*_ and *P*_*obs*_ as robust estimates of the actual species richness of the assemblage. If *A*_0_ and *P*_0_ are not zero they estimate the minimum number of undetected animal and plant species that can be expected with a sufficiently large number of replicates, taken from the same assemblage/locality by the same methods in the same time period. We can use extrapolation methods (Colwell, Dunn & Harris, 2012) to estimate how many additional replicate surveys it would take to reach a specified proportion *g* of *A*_*est*_ and *P*_*est*_.

## The interactions

We are then faced with assessing the sampling of interactions *I*. Table 1 summarizes the main components and targets for estimation of interaction richness. In contrast with traditional species diversity estimates, sampling networks has the paradox that despite the potentially interacting species being present in the sampled assemblage (i.e., included in the *A*_*obs*_ and *P*_*obs*_ species lists), some of their pairwise interactions are impossible to be recorded. The reason is forbidden links. Independently of whether we sample full communities or subset communities we face a problem: some of the interactions that we can visualize in the empty adjacency matrix Δ will simply not occur. With a total of *A*_*obs*_*P*_*obs*_ “potential” interactions (eventually augmented to *A*_*est*_*P*_*est*_ in case we have undetected species), a fraction of them are impossible to record, because they are forbidden (Jordano, Bascompte & Olesen, 2003; Olesen *et al.*, 2011).

**Table 1.** A taxonomy of link types for ecological interactions (Olesen *et al.* 2011). *A*, number of animal species; *P*, number of plant species; *I*, number of observed links; *C* = 100*I/*(*AP*), connectance; *F L*, number of forbidden links; and *M L*, number of missing links. As natural scientists, our ultimate goal is to eliminate *M L* from the equation *F L* = *AP − I − M L*, which probably is not feasible given logistic sampling limitations. When we, during our study, estimate *M L* to be negligible, we cease observing and estimate *I* and *F L*.

| Link type | Formulation | Definition |
| --- | --- | --- |
| Potential links | $I_{max} = A_{obs}P_{obs}$ | Size of observed network matrix, i.e. maximum number of potentially observable interactions; $A_{obs}$ and $P_{obs}$, numbers of interacting animal and plant species, respectively.<br>These might be below the real numbers of animal and plant species, $A_{est}$ and $P_{est}$. |
| Observed links | $I_{obs}$ | Total number of observed links in the network given a sufficient sampling effort. Number of ones in the adjacency matrix. |
| True links | $I_{est}$ | Total number of links in the network given a sufficient sampling effort; expected for the augmented $A_{est}P_{est}$ matrix. |
| Unobserved links | $UL = I_{max} - I_{obs}$ | Number of zeroes in the adjacency matrix. |
| True unobserved links | $UL* = I_{max} - I_{obs}$ | Number of zeroes in the augmented adjacency matrix that, eventually, includes unobserved species. |
| Forbidden links | $FL$ | Number of links, which remain unobserved because of linkage constraints, irrespectively of sufficient sampling effort. |
| Observed Missing links | $ML = A_{obs}P_{obs} - I_{obs} - FL$ | Number of links, which may exist in nature but need more sampling effort and/or additional sampling methods to be observed. |
| True Missing links | $ML* = A_{est}P_{est} - I_{est} - FL$ | Number of links, which may exist in nature but need more sampling effort and/or additional sampling methods to be observed.<br>Augments $ML$ for the $A_{est}P_{est}$ matrix. |

Our goal is to estimate the true number of non-null *AP* interactions, including interactions that actually occur but have not been observed (*I*_0_) from the replicated incidence frequencies of interaction types: *I*_*est*_ = *I*_*obs*_ + *I*_0_. Note that *I*_0_ estimates the minimum number of undetected plant-animal interactions that can be expected with a sufficiently large number of replicates, taken from the same assemblage/locality by the same methods in the same time period. Therefore we have two types of non-obsereved links: *U L\** and *U L*, corresponding to the real assemblage species richness and to the observed assemblage species richness, respectively (Table 1).

Forbidden links are non-occurrences of pairwise interactions that can be accounted for by biological constraints, such as spatio-temporal uncoupling (Jordano, 1987), size or reward mismatching, foraging constraints (e.g., accessibility) (Moré *et al.*, 2012), and physiological-biochemical constraints (Jordano, 1987). We still have extremely reduced information about the frequency of forbidden links in natural communities (Jordano, Bascompte & Olesen, 2003; Stang *et al.*, 2009; Vázquez, Chacoff & Cagnolo, 2009; Olesen *et al.*, 2011; Ibanez, 2012; Maruyama *et al.*, 2014; Vizentin-Bugoni, Maruyama & Sazima, 2014) (Table 1). Forbidden links are thus represented as structural zeroes in the interaction matrix, i.e., matrix cells that cannot get a non-zero value.

We might expect different types of *F L* to occupy different parts of the Δ matrix, with missing cells due to phenological uncoupling, *F L_P_*, largely distributed in the lower-right half Δ matrix and actually missed links *M L* distributed in its central part (Olesen *et al.*, 2010). Yet, most of these aspects remain understudied. Therefore, we need to account for the frequency of these structural zeros in our matrix before proceeding. For example, most measurements of connectance *C* = *I/*(*AP*) implicitly ignore the fact that by taking the full product *AP* in the denominator they are underestimating the actual connectance value, i.e., the fraction of actual interactions *I* relative to the *biologically possible* ones, not to the total maximum *I*_*max*_ = *AP*.

Our main problem then turns to estimate the number of true missed links, i.e., those that can’t be accounted for by biological constraints and that might suggest undersampling. Thus, the sampling of interactions in nature, as the sampling of species, is a cumulative process. In our analysis, we are not re-sampling individuals, but interactions, so we made interaction-based accumulation curves. If an interaction-based curve suggests a robust sampling, it does mean that no new interactions are likely to be recorded, irrespectively of the species, as it is a whole-network sampling approach (N. Gotelli, pers.com.). We add new, distinct, interactions recorded as we increase sampling effort (Fig. 2). We can obtain an Interaction Accumulation Curve (*IAC*) analogous to a Species Curve (*SAC*) (see Supplementary Online Material): the observed number of distinct pairwise interactions in a survey or collection as a function of the accumulated number of observations or samples (Colwell, 2009).

**Figure 2.**
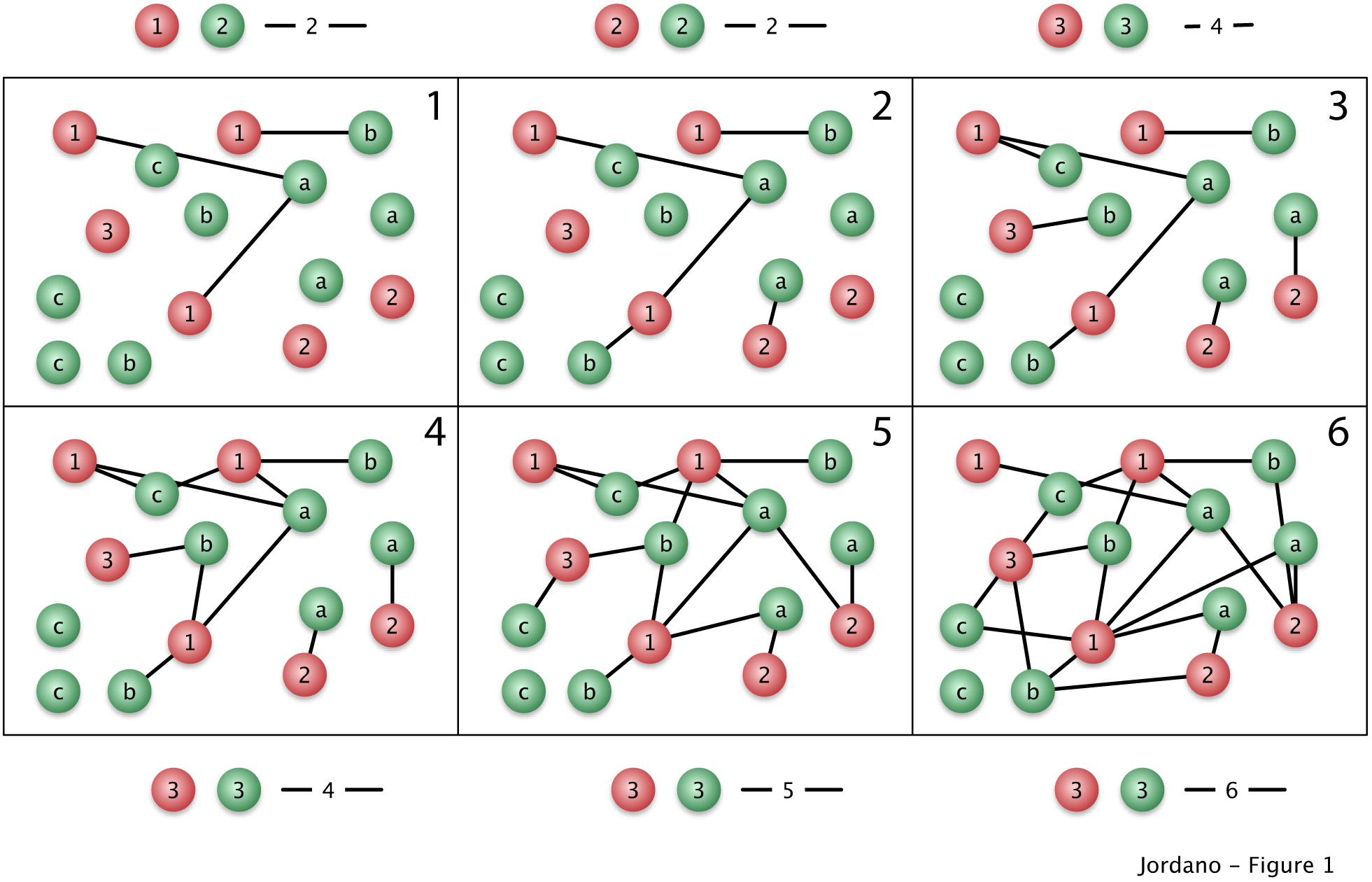
Sampling species interactions in natural communities. Suppose an assemblage with *A* = 3 animal species (red, species 1−3 with three, two, and 1 individuals, respectively) and *P* = 3 plant species (green, species a-c with three individuals each) (colored balls), sampled with increasing effort in steps 1 to 6 (panels). In Step 1 we record animal species 1 and plant species 1 and 2 with a total of three interactions (black lines) represented as two distinct interactions: 1 *− a* and 1 *− b*. As we advance our sampling (panels 1 to 6, illustrating e.g., additional sampling days) we record new distinct interactions. Note that we actually sample and record interactions among individuals, yet we pool the data across species to get a species by species interaction matrix. Few network analyses have been carried out on individual data(Dupont *et al.*, 2014).

## Empirical data on Forbidden Links

Adjacency matrices are frequently sparse, i.e., they are densely populated with zeroes, with a fraction of them being structural (unobservable interactions) (Bascompte & Jordano, 2014). Thus, it would be a serious interpretation error to attribute the sparseness of adjacency matrices for bipartite networks to undersampling. The actual typology of link types in ecological interaction networks is thus more complex than just the two categories of observed and unobserved interactions (Table 1). Unobserved interactions are represented by zeroes and belong to two categories. Missing interactions may actually exist but require additional sampling or a variety of methods to be observed. Forbidden links, on the other hand, arise due to biological constraints limiting interactions and remain unobservable in nature, irrespectively of sampling effort (Table 1). Forbidden links *F L* may actually account for a relatively large fraction of unobserved interactions *U L* when sampling taxonomically-restricted subnetworks (e.g., plant-hummingbird pollination networks) (Table 1). Phenological uncoupling is also prevalent in most networks, and may add up to explain ca. 25–40% of the forbidden links, especially in highly seasonal habitats, and up to 20% when estimated relative to the total number of unobserved interactions (Table 2). In any case, we might expect that a fraction of the missing links *M L* would be eventually explained by further biological reasons, depending on the knowledge of natural details of the particular systems. Our goal as naturalists would be to reduce the fraction of *U L* which remain as missing links; to this end we might search for additional biological constraints or increase sampling effort. For instance, habitat use patterns by hummingbirds in the Arima Valley network (Table 2; Snow & Snow, 1972) impose a marked pattern of microhabitat mismatches causing up to 44.5% of the forbidden links. A myriad of biological causes beyond those included as *F L* in Table 2 may contribute explanations for *U L*: limits of color perception and or partial preferences, presence of secondary metabolites in fruit pulp and leaves, toxins and combinations of monosaccharides in nectar, etc. For example, aside from *F L*, some pairwise interactions may simply have an asymptotically-zero probability of interspecific encounter between the partner species, if they are very rare. However, it is surprising that just the limited set of forbidden link types considered in Table 1 explain between 24.6–77.2% of the unobserved links. Notably, the Arima Valley, Santa Virgnia, and Hato Ratón networks have *>* 60% of the unobserved links explained, which might be related to the fact that they are subnetworks (Arima Valley, Santa Virgínia) or relatively small networks (Hato Ratón). All this means that empirical networks may have sizable fractions of structural zeroes. Ignoring this biological fact may contribute to wrongly inferring undersampling of interactions in real-world assemblages.

**Table 2.**
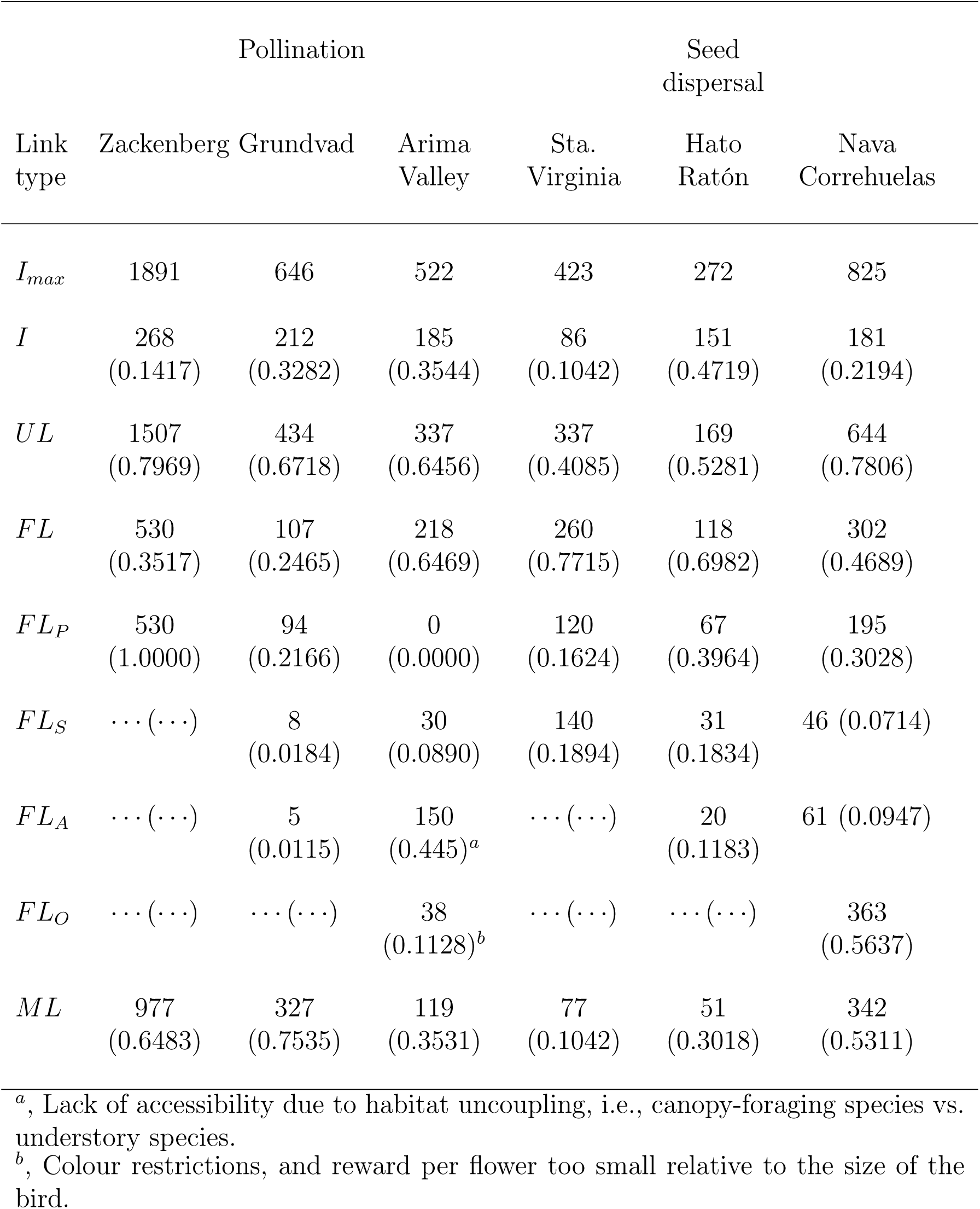
Frequencies of different type of forbidden links in natural plant-animal interaction assemblages. *AP*, maximum potential links, *I*_*max*_; *I*, number of observed links; *U L*, number of unobserved links; *F L*, number of forbidden links; *F L_P_*, phenology; *F L_S_*, size restrictions; *F L_A_*, accessibility; *F L_O_*, other types of restrictions; *M L*, unknown causes (missing links). Relative frequencies (in parentheses) calculated over *I*_*max*_ = *AP* for *I*, *M L*, and *F L*; for all forbidden links types, calculated over *F L*. References, from left to right: Olesen *et al.* 2008; Olesen & Myrthue unpubl.; Snow & Snow 1972 and Jordano *et al.* 2006; Vizentin-Bugoni *et al.* 2014; Jordano *et al.* 2009; Olesen *et al.* 2011.

To sum up, two elements of inference are required in the analysis of unobserved interactions in ecological interaction networks: first, detailed natural history information on the participant species that allows the inference of biological constraints imposing forbidden links, so that structural zeroes can by identified in the adjacency matrix. Second, a critical analysis of sampling robustness and a robust estimate of the actual fraction of missing links, *M*, resulting in a robust estimate of *I*. In the next sections I explore these elements of inference, using *IACs* to assess the robustness of interaction sampling.

## Asymptotic diversity estimates

Let’s assume a sampling of the diversity in a specific locality, over relatively homogeneous landscape where we aim at determining the number of species present for a particular group of organisms. To do that we carry out transects or plot samplings across the landscape or use any other type of direct or indirect recording method, adequately replicated so we obtain a number of samples. Briefly, *S*_*obs*_ is the total number of species observed in a sample, or in a set of samples. *S*_*est*_ is the estimated number of species in the community represented by the sample, or by the set of samples, where *est* indicates an estimator. With abundance data, let *S*_*k*_ be the number of species each represented by exactly *k* individuals in a single sample. Thus, *S*_0_ is the number of undetected species (species present in the community but not included in the sample), *S*_1_ is the number of singleton species (represented by just one individual), *S*_2_ is the number of doubleton species (species with two individuals), etc. The total number of individuals in the sample would be:

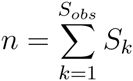

A frequently used asymptotic, bias corrected, non-parametric estimator is *S*_*Chao*1_ (Hortal, Borges & Gaspar, 2006; Chao, 2005; Colwell, 2013):

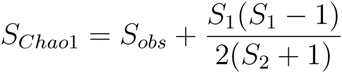

Another frequently used alternative is the Chao2 estimator, *S*_*Chao*2_ (Gotelli & Colwell, 2001), which has been reported to have a limited bias for small sample sizes (Colwell & Coddington, 1994; Chao, 2005). Instead of using counts it uses incidence frequencies (*Q*_*k*_) among samples (number of species present in just one sample, in two samples, etc.):

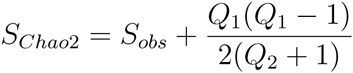

A plot of the cumulative number of species recorded, *S*_*n*_, as a function of some measure of sampling effort (say, *n* samples taken) yields the species accumulation curve (SAC) or collector’s curve (Colwell & Coddington, 1994). Similarly, interaction accumulation curves (IAC), analogous to SACs, can be used to assess the robustness of interactions sampling for plant-animal community datasets (Jordano, 1987; Jordano, Vázquez & Bascompte, 2009; Olesen *et al.*, 2011), as discussed in the next section.

## Assessing sampling effort when recording interactions

The basic method we can propose to estimate sampling effort and explicitly show the analogues with rarefaction analysis in biodiversity research is to vectorize the interaction matrix *AP* so that we get a vector of all the potential pairwise interactions (*I*_*max*_, Table 1) that can occur in the observed assemblage with *A*_*obs*_ animal species and *P*_*obs*_ plant species. The new “species” we aim to sample are the pairwise interactions (Table 3). So, if we have in our community *Turdus merula* (*T m*) and *Rosa canina* (*Rc*) and *Prunus mahaleb* (*P m*), our problem will be to sample 2 new “species”: *T m - Rc* and *T m - P m*. In general, if we have *A* = 1*…i*, animal species and *P* = 1*…j* plant species (assuming a complete list of species in the assemblage), we’ll have a vector of “new” species to sample: *A*_1_*P*_1_, *A*_1_*P*_2_,*… A*_2_*P*_1_, *A*_2_*P*_2_,*… A_i_P_j_*. We can represent the successive samples where we can potentially get records of these interactions in a matrix with the vectorized interaction matrix and columns representing the successive samples we take (Table 3). This is simply a vectorized version of the interaction matrix. This is analogous to a biodiversity sampling matrix with species as rows and sampling units (e.g., quadrats) as columns (Jordano, Vázquez & Bascompte, 2009). The package *EstimateS* (Colwell, 2013) includes a complete set of functions for estimating the mean IAC and its unconditional standard deviation from random permutations of the data, or subsampling with-out replacement (Gotelli & Colwell, 2001) and the asymptotic estimators for the expected number of distinct pairwise interactions included in a given reference sample of interaction records (see also the specaccum function in library vegan of the R Package)(R Development Core Team, 2010; Jordano, Vázquez & Bascompte, 2009; Olesen *et al.*, 2011). In particular, we may take advantage of replicated incidence data, as it takes account of heterogeneity among samples (days, censuses, etc.; R.K Colwell, pers. comm.) (see also Colwell, Mao & Chang, 2004; Colwell, Dunn & Harris, 2012; Chao *et al.*, 2014).

**Table 3.** A vectorized interaction matrix.

| Interaction | Sample 1 | Sample 2 | Sample 3 | ... | Sample $i$ |
| --- | --- | --- | --- | --- | --- |
| A1 - P2 | 12 | 2 | 0 | ... | 6 |
| A1 - P2 | 0 | 0 | 0 | ... | 1 |
| ... | ... | ... | ... | ... | ... |
| A5 - P3 | 5 | 0 | 1 | ... | 18 |
| A5 - P4 | 1 | 0 | 1 | ... | 3 |
| ... | ... | ... | ... | ... | ... |
| A<sub>i</sub> - P<sub>i</sub> | 1 | 0 | 1 | ... | 2 |

**Table 4.** Sampling statistics for three plant-animal interaction networks (Olesen *et al.* 2011). Symbols as in Table 1; *N*, number of records; *Chao*1 and *ACE* are asymptotic estimators for the number of distinct pairwise interactions *I* (Hortal *et al.* 2006), and their standard errors; *C*, sample coverage for rare interactions (Chao & Jost 2012). Scaled asymptotic estimators and their confidence intervals (*CI*) were calculated by weighting *Chao*1 and *ACE* with the observed frequencies of forbidden links.

|  | Hato Ratón | Nava Correhuelas | Zackenberg |
| --- | --- | --- | --- |
| $A$ | 17 | 33 | 65 |
| $P$ | 16 | 25 | 31 |
| $I_{max}$ | 272 | 825 | 1891 |
| $N$ | 3340 | 8378 | 1245 |
| $I$ | 151 | 181 | 268 |
| $C$ | 0.917 | 0.886 | 0.707 |
| $Chao1$ | $263.1 \pm 70.9$ | $231.4 \pm 14.2$ | $509.6 \pm 54.7$ |
| $ACE$ | $240.3 \pm 8.9$ | $241.3 \pm 7.9$ | $566.1 \pm 14.8$ |
| % *unobserved*<sup>a</sup> | 8.33 | 15.38 | 47.80 |

<sup>a</sup>, estimated with library Jade (R Core Development Team 2010, Chao *et al.* 2015)

In this way we effectively extend sampling theory developed for species diversity to the sampling of ecological interactions. Yet future theoretical work will be needed to formally assess the similarities and differences in the two approaches and developing biologically meaningful null models of expected interaction richness with added sampling effort.

Diversity-accumulation analysis (Magurran, 1988; Hortal, Borges & Gaspar, 2006) comes up immediately with this type of dataset. This procedure plots the accumulation curve for the expected number of distinct pairwise interactions recorded with increasing sampling effort (Jordano, Vázquez & Bascompte, 2009; Olesen *et al.*, 2011). Asymptotic estimates of interaction richness and its associated standard errors and confidence intervals can thus be obtained (Hortal, Borges & Gaspar, 2006) (see Supplementary Online Material). It should be noted that the asymptotic estimate of interaction richness explicitly ignores the fact that, due to forbidden links, a number of pairwise interactions among the *I*_*max*_ number specified in the adjacency matrix Δ cannot be recorded, irrespective of sampling effort.

We may expect undersampling specially in moderate to large sized networks with multiple modules (i.e., species subsets requiring different sampling strategies) (Jordano, 1987; Olesen *et al.*, 2011; Chacoff *et al.*, 2012); adequate sampling may be feasible when interaction subwebs are studied (Olesen *et al.*, 2011; Vizentin-Bugoni, Maruyama & Sazima, 2014), typically with more homogeneous subsets of species (e.g., bumblebee-pollinated flowers). In any case the sparseness of the Δ matrix is by no means an indication of undersampling whenever the issue of structural zeroes in the interaction matrices is effectively incorporated in the estimates.

For example, mixture models incorporating detectabilities have been proposed to effectively account for rare species (Mao & Colwell, 2005). In an analogous line, mixture models could be extended to samples of pairwise interactions, also with specific detectability values. These detection rate/odds could be variable among groups of interactions, depending on their specific detectability. For example, detectability of flower-pollinator interactions involving bumblebees could have a higher detectability than flower-pollinator pairwise interactions involving, say, nitidulid beetles. These more homogeneous groupings of pairwise interactions within a network define modules (Bascompte & Jordano, 2014), so we might expect that interactions of a given module (e.g., plants and their hummingbird pollinators; Fig. 1a) may share similar detectability values, in an analogous way to species groups receiving homogeneous detectability values in mixture models (Mao & Colwell, 2005). In its simplest form, this would result in a sample with multiple pairwise interactions detected, in which the number of interaction events recorded for each distinct interaction found in the sample is recorded (i.e., a column vector in Table 3, corresponding to, say, a sampling day). The number of interactions recorded for the *i*_*th*_ pairwise interaction (i.e., *A*_*i*_*P*_*j*_ in Table 3), *Y*_*i*_ could be treated as a Poisson random variable with a mean parameter *λ*_*i*_, its detection rate. Mixture models (Mao & Colwell, 2005) include estimates for abundance-based data (their analogs in interaction sampling would be weighted data), where *Y*_*i*_ is a Poisson random variable with detection rate *λ*_*i*_. This is combined with the incidence-based model, where *Y*_*i*_ is a binomial random variable (their analogous in interaction sampling would be presence/absence records of interactions) with detection odds *λ*_*i*_. Let *T* be the number of samples in an incidence-based data set. A Poisson/binomial density can be written as (Mao & Colwell, 2005):

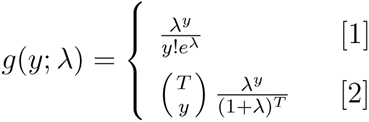

where [1] corresponds to a weighted network, and [2] to a qualitative network.

The detection rates *λ*_*i*_ depend on the relative abundances *ϕ*_*i*_ of the interactions, the probability of a pairwise interaction being detected when it is present, and the sample size (the number of interactions recorded), which, in turn, is a function of the sampling effort. Unfortunately, no specific sampling model has been developed along these lines for species interactions and their characteristic features. For example, a complication factor might be that interaction abundances, *ϕ*_*i*_, in real assemblages are a function of the abundances of interacting species that determine interspecific encounter rates; yet they also depend on biological factors that ultimately determine if the interaction occurs when the partner species are present. For example, *λ*_*i*_ should be set to zero for all *F L*. It its simplest form, *ϕ*_*i*_ could be estimated from just the product of partner species abundances, an approach recently used as a null model to assess the role of biological constraints in generating forbidden links and explaining interaction patterns (Vizentin-Bugoni, Maruyama & Sazima, 2014). Yet more complex models (e.g., Wells & O’hara 2012) should incorporate not only interspecific encounter probabilities, but also interaction detectabilities, phenotypic matching and incidence of forbidden links. Mixture models are certainly complex and for most situations of evaluating sampling effort better alternatives include the simpler incidence-based rarefaction and extrapolation (Colwell, Dunn & Harris, 2012; Chao *et al.*, 2014).

## The *real* missing links

Given that a fraction of unobserved interactions can be accounted for by for-bidden links, what about the remaining missing interactions? We have already discussed that some of these could still be related to unaccounted constraints, and still others would be certainly attributable to insufficient sampling. Would this always be the case? Multispecific assemblages of distinct taxonomic relatedness, whose interactions can be represented as bipartite networks (e.g., host-parasite, plant-animal mutualisms, plant-herbivore interactions-with two distinct sets of unrelated higher taxa), are shaped by interspecific encounters among individuals of the partner species (Fig. 2). A crucial ecological aspect limiting these inter-actions is the probability of interspecific encounter, i.e., the probability that two individuals of the partner species actually encounter each other in nature.

Given log-normally distributed abundances of the two species groups, the expected probabilities of interspecific encounter (*P IE*) would be simply the product of the two lognormal distributions. Thus, we might expect that for low *P IE* values, pairwise interactions would be either extremely difficult to sample, or just simply not occurring in nature. Consider the Nava de las Correhuelas interaction web (NCH, Table 2), with *A* = 36, *P* = 25, *I* = 181, and almost half of the unobserved interactions not accounted for by forbidden links, thus *M* = 53.1%. Given the robust sampling of this network (Jordano, Vázquez & Bascompte, 2009), a sizable fraction of these possible but missing links would be simply not occurring in nature, most likely by extremely low *P IE*, in fact asymptotically zero. Given the vectorized list of pairwise interactions for NCH, I computed the *P IE* values for each one by multiplying element-wise the two species abundance distributions. The *P IE_max_* = 0.0597, being a neutral estimate, based on the assumption that interactions occur in proportion to the species-specific local abundances. With *P IE_median_* < 1.4 10^−4^ we may safely expect (note the quantile estimate *Q*_75%_ =3.27 10^−4^) that a sizable fraction of these missing interactions may not occur according to this neutral expectation (Jordano, 1987; Olesen *et al.*, 2011) (neutral forbidden links, *sensu* Canard *et al.*, 2012).

When we consider the vectorized interaction matrix, enumerating all pairwise interactions for the *AP* combinations, the expected probabilities of finding a given interaction can be estimated with a Good-Turing approximation (Good, 1953). The technique, developed by Alan Turing and I.J. Good with applications to linguistics and word analysis (Gale & Sampson, 1995) has been recently extended in novel ways for ecological analyses (Chao *et al.*, 2015). It estimates the probability of recording an interaction of a hitherto unseen pair of partners, given a set of past records of interactions between other species pairs. Let a sample of *N* interactions so that *n*_*r*_ distinct pairwise interactions have exactly *r* records. All Good-Turing estimators obtain the underlying frequencies of events as:

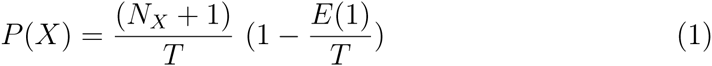

where *X* is the pairwise interaction, *N*_*X*_ is the number of times interaction *X* is recorded, *T* is the sample size (number of distinct interactions recorded) and *E*(1) is an estimate of how many different interactions were recorded exactly once. Strictly speaking Equation (1) gives the probability that the next interaction type recorded will be *X*, after sampling a given assemblage of interacting species. In other words, we scale down the maximum-likelihood estimator 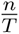 by a factor of 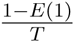. This reduces all the probabilities for interactions we have recorded, and makes room for interactions we haven’t seen. If we sum over the interactions we have seen, then the sum of *P* (*X*) is 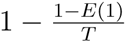. Because probabilities sum to one, we have the left-over probability of 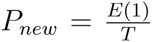 of seeing something new, where new means that we sample a new pairwise interaction. Note, however, that Good-Turing estimators, the traditional asymptotic estimators, do not account in our case for the forbidden interactions.

## Discussion

Recent work has inferred that most data available for interaction networks are incomplete due to undersampling, resulting in a variety of biased parameters and network patterns (Chacoff *et al.*, 2012). It is important to note, however, that in practice, many surveyed networks to date have been subnets of much larger networks. This is also true for protein interaction, gene regulation, and metabolic networks, where only a subset of the molecular entities in a cell have been sampled (Stumpf, Wiuf & May, 2005). Despite recent attempts to document whole ecosystem meta-networks (Pocock, Evans & Memmott, 2012), it is likely that most ecological interaction networks will illustrate just major ecosystem compartments. Due to their high generalization, high temporal and spatial turnover, and high complexity of association patterns, adequate sampling of ecological interaction networks is challenging and requires extremely large sampling effort. Undersampling of ecological networks may originate from the analysis of assemblage subsets (e.g., taxonomically or functionally defined), and/or from logistically-limited sampling effort. It is extremely hard to robustly sample the set of biotic interactions even for relatively simple, species-poor assemblages; thus, we need to assess how robust is the characterization of the adjacency matrix Δ. Concluding that an ecological network dataset is undersampled just by its sparseness would be unrealistic. The reason stems from a biological fact: a sizeable fraction of the maximum, potential links that can be recorded among two distinct sets of species is simply unobservable, irrespective of sampling effort (Jordano, 1987). In addition, sampling effort needs to be explicitly gauged because of its potential influence on parameter estimates for the network.

Missing links are a characteristic feature of all plant-animal interaction networks, and likely pervade other ecological interactions. Important natural history details explain a fraction of them, resulting in unrealizable interactions (i.e., forbidden interactions) that define structural zeroes in the interaction matrices and contribute to their extreme sparseness. Sampling interactions is a way to monitor biodiversity beyond the simple enumeration of component species and to develop efficient and robust inventories of functional interactions. Yet no sampling theory for interactions is available. Some key components of this sampling are analogous to species sampling and traditional biodiversity inventories; however, there are important differences. Focusing just on the realized interactions or treating missing interactions as the expected unique result of sampling bias would miss important components to understand how mutualisms coevolve within complex webs of interdependence among species.

Contrary to species inventories, a sizable fraction of non-observed pairwise interactions cannot be sampled, due to biological constraints that forbid their occurrence. Moreover, recent implementations of inference methods for unobserved species (Chao *et al.*, 2015) or for individual-based data (Wells & O’Hara, 2012) can be combined with the forbidden link approach. They do not account either for the existence of these ecological constraints, but can help in estimating their relative importance, simply by the difference between the asymptotic estimate of interaction richness *in a robustly-sampled* assemblage and the maximum richness *I*_*max*_ of interactions.

Ecological interactions provide the wireframe supporting the lives of species, and they also embed crucial ecosystem functions which are fundamental for supporting the Earth system. We still have a limited knowledge of the biodiversity of ecological interactions, and they are being lost (extinct) at a very fast pace, frequently preceding species extinctions (Valiente-Banuet *et al.*, 2014). We urgently need robust techniques to assess the completeness of ecological interactions networks because this knowledge will allow the identification of the minimal components of their ecological complexity that need to be restored to rebuild functional ecosystems after perturbations.

## Acknowledgements

I am indebted to Jens M. Olesen, Alfredo Valido, Jordi Bascompte, Thomas Lewinshon, John N. Thompson, Nick Gotelli, Carsten Dormann, and Paulo R. Guimaraes Jr. for useful and thoughtful discussion at different stages of this manuscript. Jeferson Vizentin-Bugoni kindly helped with the Sta Virgínia data. Jens M. Olesen kindly made available the Grundvad dataset; together with Robert K. Colwell, Néstor Pérez-Méndez, JuanPe González-Varo, and Paco Rodríguez provided most useful comments to a final version of the ms. Robert Colwell shared a number of crucial suggestions that clarified my vision of sampling ecological interactions. The study was supported by a Junta de Andalucía Excellence Grant (RNM–5731), as well as a Severo Ochoa Excellence Award from the Ministerio de Economía y Competitividad (SEV–2012–0262). The Agencia de Medio Ambiente, Junta de Andalucía, provided generous facilities that made possible my long-term field work in different natural parks.

## Data accessiblity

This review does not use new raw data, but includes some re-analyses of previously published material. All the original data supporting the paper, R code, supplementary figures, and summaries of analytical protocols is available at the author’s GitHub repository (

~~~
https://github.com/pedroj/MS_Network-Sampling
~~~

), with DOI: 

~~~
10.5281/zenodo.29437
~~~

.

## References

Banasek-Richter, C., Cattin, M. & Bersier, L. (2004) Sampling effects and the robustness of quantitative and qualitative food-web descriptors. Journal of Theoretical Biology, 226, 23–32.

Bascompte, J. & Jordano, P. (2014) Mutualistic networks. Monographs in Population Biology, No. 53. Princeton University Press, Princeton, NJ.

Bersier, L., Banasek-Richter, C. & Cattin, M. (2002) Quantitative descriptors of food-web matrices. Ecology, 83, 2394–2407.

Bluthgen, N. (2010) Why network analysis is often disconnected from community ecology: A critique and an ecologist’s guide. Basic And Applied Ecology, 11, 185–195.

Bosch, J., Martín González, A.M., Rodrigo, A. & Navarro, D. (2009) Plantpollinator networks: adding the pollinator’s perspective. Ecology Letters, 12, 409–419.

Brose, U., Martinez, N. & Williams, R. (2003) Estimating species richness: Sensitivity to sample coverage and insensitivity to spatial patterns. Ecology, 84, 2364–2377.

Bunge, J. & Fitzpatrick, M. (1993) Estimating the number of species: a review. Journal of the American Statistical Association, 88, 364–373.

Canard, E., Mouquet, N., Marescot, L., Gaston, K.J., Gravel, D. & Mouillot, D. (2012) Emergence of structural patterns in neutral trophic networks. PLoS ONE, 7, e38295.

Chacoff, N.P., Vazquez, D.P., Lomascolo, S.B., Stevani, E.L., Dorado, J. & Padrón, B. (2012) Evaluating sampling completeness in a desert plant-pollinator network. Journal of Animal Ecology, 81, 190–200.

Chao, A. (2005) Species richness estimation. Encyclopedia of Statistical Sciences, pp. 7909–7916. Oxford University Press, New York, USA.

Chao, A., Gotelli, N.J., Hsieh, T.C., Sander, E.L., Ma, K.H., Colwell, R.K. & Ellison, A.M. (2014) Rarefaction and extrapolation with Hill numbers: a framework for sampling and estimation in species diversity studies. Ecological Monographs, 84, 45–67.

Chao, A., Hsieh, T.C., Chazdon, R.L., Colwell, R.K. & Gotelli, N.J. (2015) Unveiling the species-rank abundance distribution by generalizing the Good-Turing sample coverage theory. Ecology, 96, 1189–1201.

Coddington, J.A., Agnarsson, I., Miller, J.A., Kuntner, M. & Hormiga, G. (2009) Undersampling bias: the null hypothesis for singleton species in tropical arthropod surveys. Journal of Animal Ecology, 78, 573–584.

Cohen, J.E. (1978) Food webs and niche space. Princeton University Press, Princeton, New Jersey, US.

Cohen, J.E., Beaver, R.A., Cousins, S.H., DeAngelis, D.L., Goldwasser, L., Heong, K.L., Holt, R.D., Kohn, A.J., Lawton, J.H., Martinez, N., O’Malley, R., Page, L.M., Patten, B.C., Pimm, S.L., Polis, G., Rejmanek, M., Schoener, T.W., Schenly, K., Sprules, W.G., Teal, J.M., Ulanowicz, R., Warren, P.H., Wilbur, H.M. & Yodis, P. (1993) Improving food webs. Ecology, 74, 252–258.

Colwell, R. & Coddington, J. (1994) Estimating terrestrial biodiversity through extrapolation. Philosophical Transactions Of The Royal Society Of London Series B-Biological Sciences, 345, 101–118.

Colwell, R.K. (2009) Biodiversity: concepts, patterns, and measurement. The Princeton Guide to Ecology (ed. S.A. Levin), pp. 257–263. Princeton University Press, Princeton.

Colwell, R.K. (2013) EstimateS: Biodiversity Estimation. -, pp. 1–33.

Colwell, R.K., Dunn, R.R. & Harris, N.C. (2012) Coextinction and persistence of dependent species in a changing world. Annual Review of Ecology Evolution and Systematics, 43, 183–203.

Colwell, R.K., Mao, C.X. & Chang, J. (2004) Interpolating, extrapolating, and comparing incidence-based species accumulation curves. Ecology, 85, 2717–2727.

Dorado, J., Vazquez, D.P., Stevani, E.L. & Chacoff, N.P. (2011) Rareness and specialization in plant-pollinator networks. Ecology, 92, 19–25.

Dormann, C.F., Frund, J., Bluthgen, N. & Gruber, B. (2009) Indices, graphs and null models: Analyzing bipartite ecological networks. Open Ecology Journal, 2, 7–24.

Dupont, Y.L., Trøjelsgaard, K. & Olesen, J.M. (2011) Scaling down from species to individuals: a flower–visitation network between individual honeybees and thistle plants. Oikos, 120, 170–177.

Dupont, Y.L., Trøjelsgaard, K., Hagen, M., Henriksen, M.V., Olesen, J.M., Pedersen, N.M.E. & Kissling, W.D. (2014) Spatial structure of an individual-based plant-pollinator network. Oikos, 123, 1301–1310.

Eklöf, A., Jacob, U., Kopp, J., Bosch, J., Castro-Urgal, R., Chacoff, N.P., Dalsgaard, B., de Sassi, C., Galetti, M., Guimaraes, P.R., Lomáscolo, S.B., Martín González, A.M., Pizo, M.A., Rader, R., Rodrigo, A., Tylianakis, J.M., Vazquez, D.P. & Allesina, S. (2013) The dimensionality of ecological networks. Ecology Letters, 16, 577–583.

Elberling, H. & Olesen, J.M. (1999) The structure of a high latitude plant-flower visitor system: the dominance of flies. Ecography, 22, 314–323.

Frund, J., McCann, K.S. & Williams, N.M. (2015) Sampling bias is a challenge for quantifying specialization and network structure: lessons from a quantitative niche model. Oikos, pp. n/a–n/a

Gale, W.A. & Sampson, G. (1995) Good-Turing frequency estimation without tears. Journal of Quantitative Linguistics, 2, 217–237.

Gibson, R.H., Knott, B., Eberlein, T. & Memmott, J. (2011) Sampling method influences the structure of plant–pollinator networks. Oikos, 120, 822–831.

González-Varo, J.P., Arroyo, J.M. & Jordano, P. (2014) Who dispersed the seeds? The use of DNA barcoding in frugivory and seed dispersal studies. Methods in Ecology and Evolution, 5, 806–814.

Good, I.J. (1953) The population frequencies of species and the estimation of population parameters. Biometrika, 40, 237–264.

Gotelli, N.J. & Colwell, R.K. (2011) Estimating species richness. Biological Diversity Frontiers in Measurement and Assessment (eds. A.E. Magurran & B.J. McGill), pp. 39–54. Oxford University Press, Oxford, UK.

Gotelli, N. & Colwell, R. (2001) Quantifying biodiversity: procedures and pitfalls in the measurement and comparison of species richness. Ecology Letters, 4, 379–391.

Hortal, J., Borges, P. & Gaspar, C. (2006) Evaluating the performance of species richness estimators: sensitivity to sample grain size. Journal of Animal Ecology, 75, 274–287.

Ibanez, S. (2012) Optimizing size thresholds in a plant–pollinator interaction web: towards a mechanistic understanding of ecological networks. Oecologia, 170, 233–242.

Jordano, P. (1987) Patterns of mutualistic interactions in pollination and seed dispersal: connectance, dependence asymmetries, and coevolution. The American Naturalist, 129, 657–677.

Jordano, P., Bascompte, J. & Olesen, J. (2003) Invariant properties in coevolutionary networks of plant-animal interactions. Ecology Letters, 6, 69–81.

Jordano, P., Vázquez, D. & Bascompte, J. (2009) Redes complejas de interacciones planta—animal. Ecología y evolución de interacciones planta-animal (eds. R. Medel, R. Dirzo & R. Zamora), pp. 17–41. Editorial Universitaria, Santiago, Chile.

Jurado-Rivera, J.A., Vogler, A.P., Reid, C.A.M., Petitpierre, E. & Gomez-Zurita, J. (2009) DNA barcoding insect-host plant associations. Proceedings Of The Royal Society B-Biological Sciences, 276, 639–648.

Magurran, A. (1988) Ecological diversity and its measurement. Princeton University Press, Princeton, US.

Mao, C. & Colwell, R.K. (2005) Estimation of species richness: mixture models, the role of rare species, and inferential challenges. Ecology, 86, 1143–1153.

Martinez, N.D. (1993) Effects of resolution on food web structure. Oikos, 66, 403–412.

Martinez, N. (1991) Artifacts or attributes? Effects of resolution on food-web patterns in Little Rock Lake food web. Ecological Monographs, 61, 367–392.

Maruyama, P.K., Vizentin-Bugoni, J., Oliveira, G.M., Oliveira, P.E. & Dalsgaard, B. (2014) Morphological and spatio-temporal mismatches shape a neotropical savanna plant-hummingbird network. Biotropica, 46, 740–747.

Moré, M., Amorim, F.W., Benitez-Vieyra, S., Medina, A.M., Sazima, M. & Cocucci, A.A. (2012) Armament Imbalances: Match and Mismatch in PlantPollinator Traits of Highly Specialized Long-Spurred Orchids. PLoS ONE, 7, e41878.

Morris, R.J., Gripenberg, S., Lewis, O.T. & Roslin, T. (2013) Antagonistic interaction networks are structured independently of latitude and host guild. Ecology Letters, 17, 340–349.

Nielsen, A. & Bascompte, J. (2007) Ecological networks, nestedness and sampling effort. Journal of Ecology, 95, 1134–1141–1141.

Olesen, J.M., Bascompte, J., Dupont, Y.L., Elberling, H. & Jordano, P. (2011) Missing and forbidden links in mutualistic networks. Proceedings Of The Royal Society B-Biological Sciences, 278, 725–732.

Olesen, J.M., Dupont, Y.L., O’gorman, E., Ings, T.C., Layer, K., Melin, C.J., Trjelsgaard, K., Pichler, D.E., Rasmussen, C. & Woodward, G. (2010) From Broadstone to Zackenberg. Advances in Ecological Research, 42, 1–69.

Olesen, J. & Jordano, P. (2002) Geographic patterns in plant-pollinator mutualistic networks. Ecology, 83, 2416–2424.

Olito, C. & Fox, J.W. (2014) Species traits and abundances predict metrics of plant-pollinator network structure, but not pairwise interactions. Oikos, 124, 428–436.

Ollerton, J. & Cranmer, L. (2002) Latitudinal trends in plant-pollinator interactions: are tropical plants more specialised? Oikos, 98, 340–350.

Pereira, H.M., Ferrier, S., Walters, M., Geller, G.N., Jongman, R.H.G., Scholes, R.J., Bruford, M.W., Brummitt, N., Butchart, S.H.M., Cardoso, A.C., Coops, N., Dulloo, E., Faith, D., Freyhof, J., Gregory, R.D., Heip, C., Hoft, R., Hurtt, G., Jetz, W., Karp, D.S., Mcgeoch, M., Obura, D., Onoda, Y., Pettorelli, N., Reyers, B., Sayre, R., Scharlemann, J.P.W., Stuart, S., Turak, E., Walpole, M. & Wegmann, M. (2013) Essential biodiversity variables. Science, 339, 277–278.

Pocock, M.J.O., Evans, D.M. & Memmott, J. (2012) The Robustness and Restoration of a Network of Ecological Networks. Science, 335, 973–977.

Preston, F. (1948) The commonness, and rarity, of species. Ecology, 29, 254–283.

R Development Core Team (2010) R: A language and environment for statistical computing. R Foundation for Statistical Computing. Vienna, Austria. http://www.R-project.org, Vienna, Austria.

Rivera-Hutinel, A., Bustamante, R.O., Marín, V.H. & Medel, R. (2012) Effects of sampling completeness on the structure of plant-pollinator networks. Ecology, 93, 1593–1603.

Schleuning, M., Frund, J., Klein, A.M., Abrahamczyk, S., Alarcón, R., Albrecht, M., Andersson, G.K.S., Bazarian, S., Böhning-Gaese, K., Bommarco, R., Dalsgaard, B., Dehling, D.M., Gotlieb, A., Hagen, M., Hickler, T., Holzschuh, A., Kaiser-Bunbury, C.N., Kreft, H., Morris, R.J., Sandel, B., Sutherland, W.J., Svenning, J.C., Tscharntke, T., Watts, S., Weiner, C.N., Werner, M., Williams, N.M., Winqvist, C., Dormann, C.F. & Blüthgen, N. (2012) Specialization of mutualistic interaction networks decreases toward tropical latitudes. Current Biology, 22, 1925–1931.

Snow, B. & Snow, D. (1972) Feeding niches of hummingbirds in a Trinidad valley. Journal of Animal Ecology, 41, 471–485.

Snow, B. & Snow, D. (1988) Birds and berries. Poyser, Calton, UK.

Stang, M., Klinkhamer, P., Waser, N.M., Stang, I. & van der Meijden, E. (2009) Size-specific interaction patterns and size matching in a plant-pollinator interaction web. Annals Of Botany, 103, 1459–1469.

Strogatz, S. (2001) Exploring complex networks. Nature, 410, 268–276.

Stumpf, M.P.H., Wiuf, C. & May, R.M. (2005) Subnets of scale-free networks are not scale-free: Sampling properties of networks. Proceedings of the National Academy of Sciences USA, 102, 4221–4224.

Thébault, E. & Fontaine, C. (2010) Stability of ecological communities and the architecture of mutualistic and trophic networks. Science, 329, 853–856.

Valiente-Banuet, A., Aizen, M.A., Alcántara, J.M., Arroyo, J., Cocucci, A., Galetti, M., García, M.B., García, D., Gomez, J.M., Jordano, P., Medel, R., Navarro, L., Obeso, J.R., Oviedo, R., Ramírez, N., Rey, P.J., Traveset, A., Verdú, M. & Zamora, R. (2014) Beyond species loss: the extinction of ecological interactions in a changing world. Functional Ecology, 29, 299–307.

Vazquez, D.P., Chacoff, N.P. & Cagnolo, L. (2009) Evaluating multiple determinants of the structure of plant-animal mutualistic networks. Ecology, 90, 2039–2046.

Vazquez, D.P., Ramos-Jiliberto, R., Urbani, P. & Valdovinos, F.S. (2015) A conceptual framework for studying the strength of plant-animal mutualistic interactions. Ecology Letters, 18, 385–400.

Vazquez, D., Morris, W. & Jordano, P. (2005) Interaction frequency as a surrogate for the total effect of animal mutualists on plants. Ecology Letters, 8, 1088–1094.

Vizentin-Bugoni, J., Maruyama, P.K. & Sazima, M. (2014) Processes entangling interactions in communities: forbidden links are more important than abundance in a hummingbird-plant network. Proceedings Of The Royal Society B-Biological Sciences, 281, 20132397–20132397.

Wells, K. & O’Hara, R.B. (2012) Species interactions: estimating per-individual interaction strength and covariates before simplifying data into per-species ecological networks. Methods in Ecology and Evolution, 4, 1–8.

Wirta, H.K., Hebert, P.D.N., Kaartinen, R., Prosser, S.W., Várkonyi, G. & Roslin, T. (2014) Complementary molecular information changes our perception of food web structure. Proceedings of the National Academy of Sciences USA, 111, 1885–1890.

